# Protein S-nitrosylation of Human Cytomegalovirus pp71 inhibits its ability to limit STING antiviral responses

**DOI:** 10.1101/2020.01.08.899757

**Authors:** Masatoshi Nukui, Kathryn L. Roche, Jie Jia, Paul L. Fox, Eain A. Murphy

## Abstract

Human Cytomegalovirus (HCMV) is a ubiquitous pathogen that has co-evolved with its host and in doing so, is highly efficient in undermining antiviral responses that limit successful infections. As a result, HCMV infections are highly problematic in individuals with weakened or underdeveloped immune systems including transplant recipients and newborns. Understanding how HCMV controls the microenvironment of an infected cell so as to favor productive replication is of critical importance. To this end, we took an unbiased proteomics approach to identify the highly reversible, stress induced, post-translational modification (PTM), protein S-nitrosylation, on viral proteins to determine the biological impact on viral replication.

We identified protein S-nitrosylation of 13 viral proteins during infection of highly permissive fibroblasts. One of these proteins, pp71, is critical for efficient viral replication, as it undermines host antiviral responses, including STING activation. By exploiting site-directed mutagenesis of the specific amino acids we identified in pp71 as protein S-nitrosylated, we found this pp71 PTM diminishes its ability to undermine antiviral responses induced by the STING pathway. Our results suggest a model in which protein S-nitrosylation may function as a host response to viral infection that limits viral spread.

**IMPORTANCE:** In order for a pathogen to establish a successful infection, it must undermine the host cell responses inhibitory to the pathogen. As such, herpesviruses encode multiple viral proteins that antagonize each host antiviral response, thereby allowing for efficient viral replication. Human Cytomegalovirus encodes several factors that limit host countermeasures to infection, including pp71. We identified a previously unreported modification of pp71 residues within the protein are protein S-nitrosylated. Using site-directed mutagenesis, we mutated the specific sites of this modification thereby blocking this pp71 post-translational modification. In contexts where pp71 is not protein S-nitrosylated, host antiviral response was inhibited. The net result of this post-translational modification is to render a viral protein with diminished abilities to block host responses to infection. This novel work supports a model in which protein S-nitrosylation may be an additional mechanism in which a cell inhibits a pathogen during the course of infection.

## INTRODUCTION

Human cytomegalovirus (HCMV) is a wide-spread betaherpesvirus that has successfully established infections within the majority of the human population. The incidence of HCMV sero-positivity increases with age (1) and infections are life-long, as HCMV undergoes both lytic and latent lifecycles (2). The lytic lifecycle is highly efficient at undermining the host antiviral responses (3), whereas during the latent stage the virus remains dormant, thereby assuring immune evasion and allowing for lifelong infections (4). Primary infections are often resolved by a competent immune system and maintenance of the latent genome is often clinically inapparent. However, HCMV infections in individuals with weakened or underdeveloped immune systems often result in poor outcomes. For example, HCMV infections of neonates can cause developmental issues, including premature births, blindness, deafness, and learning disabilities with incidence rates exceeding that of congenital birth defects due to Zika virus infection (5). Equally problematic are viral reactivation events within immunosuppressed or immunodepleted patient populations, such as organ recipients and those undergoing aggressive chemotherapy. Viral reactivations within these individuals can lead to viremia, end-stage organ failure, donor organ rejection, and graft versus host disease, all resulting in poor patient prognosis (6). Consequently, HCMV is a significant economic and medical burden.

HCMV is a large DNA virus that encodes more than 200 open reading frames (ORFs) (7, 8), of which many remain uncharacterized, partly due to the lack of non-human model systems. While there are related species of cytomegaloviruses, each have evolved to restrict their infections to tissues derived only from their host species. Even with this restricted tropism, cytomegaloviruses are highly successful pathogens, as these viruses have co-evolved with their hosts (9) and have acquired mechanisms to challenge known antiviral responses (10). In many cases, cytomegaloviruses, including HCMV, not only undermine these responses but also utilize them for the virus’ own replication. One such example is the viral tegument protein, pp71, encoded by the UL82 ORF (11). pp71 is critical for efficient HCMV replication, as viruses that lack this ORF have severe growth defects (12, 13). pp71 functions by binding to both retinoblastoma protein (pRb) (14, 15) and the Death Associated Protein 6 (DAXX) (16, 17), which results in their degradation. pRb interacts with histone deacetylases and leads to the establishment of transcriptionally repressive nucleosome formation on chromatin (18). DAXX is also a potent transcriptional repressor that functions by binding to transcription factors (19). Thus, pp71-mediated binding and degradation of these two proteins permits the virus to establish a subcellular environment conducive for viral lytic replication. pp71 expression also reduces the expression of major histocompatibility complex (MHC) on the cell surface (20) and interacts with components of the innate immune cytoplasmic DNA sensing machinery, including pp71-mediated inhibition of the Stimulator of Interferon Genes (STING) pathway (21). STING, encoded by transmembrane protein 173 (TMEM173), is a potent mediator of the interferon signaling pathway (22). Activation of cytoplasmic DNA sensors results in the production of cyclic dinucleotides that serve as activators of STING. Cyclic dinucleotide bound STING results in stimulation of both NFkB and IRF3, thereby inducing the type I interferon response to invading intracellular pathogens (23). Thus, pp71’s ability to circumvent STING activation is critical for undermining the host antiviral response and promoting a proviral state.

While viruses can challenge host responses, this is not without countermeasures from the host including post-translational modifications (PTMs) of viral proteins. PTMs are responsible for altering functions, localization, and stability of proteins. The most frequently reported PTMs are phosphorylation of serine, threonine, and tyrosine; methylation, SUMOylation, and ubiquitination of lysine and arginine; acetylation of lysine; disulfide bond linkage of cysteine; and glycosylation and lipidation of several different amino acids. One modification that is highly prevalent, yet under-studied in the field of PTMs, is protein S-nitrosylation. Protein S-nitrosylation is the covalent linkage of a nitric oxide group to the reduced thiol of cysteine, thereby increasing the size and polarity of this amino acid (24). There are multiple consequences of protein S-nitrosylation, including the allosteric regulation of a protein’s enzymatic activity in response to intracellular nitric oxide levels, regulation of protein-protein interactions, and alterations in protein folding by blocking potential disulfide linkages and preventing palmitoylation of cysteine (25), all of which would serve as effective mechanisms to limit the function of viral proteins during productive infections. The reversible nature of the S-nitrosylation linkage to the thiol group makes studying this PTM difficult. S-nitrosylated reactions are reversed under reducing conditions and are often lost during standard isolation techniques utilized in protein purification and subsequent analyses (26). Thus, this PTM is significantly under-represented in the literature.

With recent advancements in whole cell proteomic analyses coupled with innovative stable labeling of S-nitrosylated cysteines, the detection of protein S-nitrosylation can now be applied to complex biological systems (27). As the stress of HCMV infections result in global and profound changes within host cells, including the stimulation of inducible nitric oxide synthase (iNOS) (28), we sought to determine the biological impact of protein S-nitrosylation following HCMV infection. Herein, we evaluated the impact of HCMV infection on the protein S-nitrosylation state on the proteome of infected fibroblasts using unbiased, whole cell proteomics. We identified specific protein S-nitrosylated PTMs for a subset of HCMV proteins, including pp71, a viral protein that limits host antiviral responses including STING activation. Additionally, mutation of the protein S-nitrosylated cysteines within pp71 to the closely related amino acid, serine resulted in increased inhibition of STING antiviral responses, suggesting protein S-nitrosylation may negatively impact the functions of viral proteins. We report that protein S-nitrosylation of pp71 is an important regulatory mechanism that controls this viral protein’s biological function. Importantly, these findings suggest that protein S-nitrosylation of viral proteins by the host cell machinery may function as a universal antiviral defense mechanism that may inhibit multiple types of viruses.

## MATERIALS AND METHODS

### Cell culture

Primary newborn human foreskin fibroblasts (NuFF-1; GlobalStem) or primary human embryonic lung fibroblasts (MRC5; ATCC) were maintained in Dulbecco’s modified Eagle medium (DMEM) supplemented with 10% fetal bovine serum (FBS) and 100 U/ml each penicillin and streptomycin. Human Embryonic Kidney 293 FT cells (293FT; ATCC) and amphomorphic phoenix cells (Phoenix-Ampho; ATCC) were maintained in DMEM supplemented with 10% newborn calf serum (NCS) and 100 U/ml each penicillin and streptomycin. Unless indicated, all cells were maintained at 37°C/5% CO_2_.

### Biotin-switch assay and mass spectrometry

NuFF-1 cells were infected with TB40/E-mCherry-UL99eGFP (29) (herein referred to as wild-type (WT) virus) at a multiplicity of infection (moi) of 3. Infected cells were harvested 96 hours post-infection (hpi), and the biotin-switch assay was conducted using S-Nitrosylated Protein Detection kit (Cayman), following the manufacturer’s instructions. Proteins were trypsinized, and biotinylated peptides were isolated by Avidin agarose resin (Pierce). The eluates were evaporated using a SpeedVac centrifuge concentrator. Dried samples were resuspended in 10 µL of 0.1% trifluoroacetic acid then purified by solid phase extraction (SPE) using C18 Ziptip (stage-tip; EMD Millipore) following the manufacture’s protocol. Next, 5 μL of the extract in 1% acetic acid was injected and the peptides eluted from the column (Dionex 15 cm x 75 μm id Acclaim Pepmap C18, 2μm, 100 Å reversed phase capillary chromatography column) by an acetonitrile/0.1% formic acid gradient at a flow rate of 0.3 μL/min and were introduced into the source of the mass spectrometer (LTQ-Obitrap Elite hybrid mass spectrometer, Finnigan). The microelectrospray ion source was operated at 2.1 kV. The digest was analyzed using the data-dependent multitask capability of the instrument, acquiring full-scan mass spectra to determine peptide molecular weights and product ion spectra to determine amino acid sequence in successive instrument scans. The data were analyzed by using all CID spectra collected in the experiment to search the human and HCMV/HHV5 reference sequence (NCBI:txid295027) with the search program, Sequest (30). Biological replicates from three independent infections of both HCMV-infected and mock samples were analyzed by mass spectrometry.

### BAC recombineering, virus production, and viral growth analyses

Based on the mass spectrometry results, the identified S-nitrosylated cysteines (C) were mutated to serine (S). S-nitrosylation-resistant mutants (pp71 single mutant, C218S and triple mutant, C34S/C94S/C218S) were generated using the BAC-derived parental virus, TB40/E-mCherry-UL99-eGFP, using the galK positive-/counter-selection scheme of BAC recombineering, described previously (31). In brief, *galK* was amplified using primers designed with 50-bp homology to the flanking region of each mutation site (Supplemental Table 1). Purified PCR products were transformed into electrocompetent SW105 *Escherichia coli* (*E. coli*) cells harboring the TB40/E-mCherry-UL99eGFP BAC genome. GalK-positive clones were screened by sequence verification. Positive recombinants were made electrocompetent and double-stranded DNA oligos in which the codon for cysteine (TGT) was replaced by the codon for serine (TCA) flanked by homology arms, were then transformed into electrocompetent SW105 *E. coli* cells that harbored the galK-positive intermediate BAC genome. Successful recombinants were then counter-selected on 2-deoxygalactose-containing plates, and the desired mutation was confirmed by Sanger sequencing.

Viral BAC DNA was isolated as described previously (32). Viral BAC DNA and the HCMV pp71-expressing plasmid, pCGN1-pp71 (1 μg) (33) were transfected into low passage MRC5 cells by electroporation, transferred to 10 cm dishes and incubated at 37°C/5% CO_2_ overnight. After 24h post-transfection, cells were washed 1x with phosphate-buffered saline (PBS) to remove dead cells and refed with fresh DMEM supplemented with 10% FBS and 100 U/ml each penicillin and streptomycin. After 7 days (d), the transfected cells were trypsonized and 1/10^th^ of the transfection was divided between 12 145 mm dishes containing confluent MRC5 cells, each cultured in 20 ml DMEM, supplemented with 10% new born calf serum (NBCS) and 100 U/ml each penicillin and streptomycin. After 100% cytopathic effect (CPE) was observed, infected cells and media were collected and transferred to centrifuge tubes. Cells were pelleted and supernatant was reserved. Virus was released from the cell pellets by resuspension in 10 ml of media and disruption by bath sonication. The medium was then cleared of cellular debris, and the supernatant was added to the reserved medium. Virus was then purified by ultracentrifugation through a 20% sorbitol cushion (20% sorbitol in 1X PBS) for 90 minutes (min) at 72,000 × *g* at 25°C. The resulting pellet was resuspended in 5mls of DMEM supplemented with 10% FBS and 1.5% bovine serum albumin (BSA) in a 1:1 ratio and aliquots were stored at −80°C following snap-freezing in liquid nitrogen. Virus stocks were titered by 50% tissue culture infectious dose (TCID_50_) on naïve NuFF-1 cells.

Newly generated recombinant S-nitrosylation-resistant viruses (pp71 single mutant, C218S and triple mutant, C34S/C94S/C218S) were compared to the parental, wild-type TB40/E-mCherry-UL99eGFP (WT) by infecting a confluent monolayer of MRC5 cells at an moi of 1 infectious unit (IU)/cell as quantified by TCID_50_ assay. Supernatants were collected every day for a week and viral titers quantified by TCID_50_ assay on naïve NUFF-1 cells.

### Generation of plasmids for transfection and transduction

Primers were generated to amplify the different pp71 ORFs from the WT and recombinant viruses for subsequent cloning into expression constructs using the primer sets (Supplemental Table 1). FLAG-epitope tagged pp71-WT or pp71-C218S were generated using the primers in Table S1 and the respective BAC constructs as templates. The PCR products were digested with EcoRI and then cloned into pCMSeGFP (Clonetech) using the EcoRI site within the multicloning site. To generate pp71-expressing MRC5 cells, pp71 WT or pp71 C218S were PCR amplified from the above BAC constructs and subsequently cloned into the pLXSN retroviral vector (Clonetech) using the EcoRI site. pCDNA-STING-HA was a generous gift from Dr. Blossom A. Damania (UNC-Chapel Hill) (34).

To generate pp71 expressing retroviruses, phoenix cells were transfected with 1µg of either pLXSN-pp71WT or pLXSN-pp71 C218S using Lipofectamine 2000 (Invitrogen) according to the manufacture’s protocol. Transfected Phoenix cells were washed, replenished with media and maintained in at 32°C/5% CO_2_ for 2 d. The retrovirus-containing media was then clarified by low-speed centrifugation and added to MRC5 cells at 70% confluency. The following day, successfully transduced MRC5 cells were selected by adding 300 µg/mL G418 (Invitrogen) to the media. This selection treatment was repeated for a total of two cell doublings of the MRC5 cells.

### Co-immunoprecipitation and western blots

For western blotting experiments, total protein was extracted in lysis buffer (35) containing 20 mM Hepes (pH 7.4), 50 mM NaCl, 1.5 mM MgCl2, 2 mM dithiothreitol (DTT), 2 mM EGTA, 10 mM NaF, 12.5 mM β-glycerophosphate, 1 mM Na3VO4, 5 mM Na-pyrophosphate, 0.2% (v/v) Triton X-100, and Complete protease inhibitors (Roche Applied Science). An equal amount of protein (20 μg) was separated by 8% SDS-PAGE, transferred to nitrocellulose/PVDF by semi-dry transfer, and then probed with the following primary antibodies: anti-pp71 (1:100; 2H10-9 (14), anti-IE2 (1:100; 3A9 (36), anti-pp65 (1:100; 8A8 (37), anti-STING (1:1000; D2P2F, Cell Signaling), anti-biotin (1:75; Cayman), anti-FLAG (1:2000; Sigma-Aldrich), anti-HA (1:1000, Sigma Aldrich) and anti-α-tubulin (1:5,000; Sigma-Aldrich). Membranes were incubated with a secondary antibody: IRDye 800CW anti-rabbit antibody (1:15000, LI-COR) or IR-Dye 680RD anti-mouse antibody (1:15000, LI-COR). Membranes were then washed with 1X tris-buffered saline (TBS), pH 7.6 supplemented with 0.05% Tween 20 (1xTBS-T), scanned using an Odyssey (LI-COR), and protein bands were visualized by Image Studio software (LI-COR).

For all affinity/immunoprecipitation (IP) experiments, cells were harvested at 4 dpi and lysed in IP lysis buffer containing 20 mM Hepes (pH 7.4), 50 mM NaCl, 1.5 mM MgCl2, 2 mM DTT, 2 mM EGTA, 10 mM NaF, 12.5 mM β-glycerophosphate, 1 mM Na_3_VO_4_, 5 mM Na-pyrophosphate, 0.2% (v/v) Triton X-100, and Complete protease inhibitors (Roche Applied Science). To confirm S-nitrosylation of pp71, NuFF-1 cells were infected at a multiplicity of 1 IU/cell with TB40/E-mCherry-UL99eGFP. After the biotin-switch assay, lysates were affinity-purified with avidin beads or immunoprecipitated with the indicated antibody. Purified lysates were separated by 8% SDS-PAGE and blotted onto Protran 0.45µm nitrocellulose membrane (Amersham). S-nitrosylated proteins were detected by using HCMV-specific antibodies (1:100), as indicated, or an antibody against biotin (1:75).

For IP, lysates were pre-cleared for 1 hour (h) with mouse immunoglobulin G (IgG) agarose or rabbit immunoglobulin G (IgG) agarose (Sigma), dependent on the species of antibody used in IP, incubated overnight with IP antibodies and then with protein A/G beads for 1 h with rotation. After incubation, beads were washed with IP lysis buffer and boiled with 4X Laemmli sample buffer. An equal amount of protein (20 μg) was separated by 8% SDS-PAGE, transferred to nitrocellulose/PVDF, and then probed with the following primary antibodies: anti-m2 FLAG (1:7,500; Sigma-Aldrich) and anti-α-tubulin (1:5,000; Sigma-Aldrich). Both were visualized using a secondary antibody conjugated to horseradish peroxidase (goat-anti-mouse HRP, 1:10,000; Jackson ImmunoResearch).

### Quantification of DNA and RNA

Total DNA was extracted by phenol:chloroform:Isoamyl purification followed by ethanol precipitation. DNA concentrations were then determined, and equal amounts of DNA were analyzed by quantitative PCR (qPCR) in triplicate using *Power* SYBR Green Master Mix (Applied Biosystems) and an Eppendorf Mastercycler RealPlex_2_ real-time PCR machine. Copy number was calculated using a standard curve generated from HCMV BAC DNA, containing eGFP and mdm2, a kind gift from Dr. Christine O’Connor (Cleveland Clinic). Primers used for qPCR are listed in Supplemental Table 1.

Total RNA was isolated using TRIzol Reagent (Invitrogen) under RNAse-free conditions, according to the manufacturer’s instructions, and then precipitated with isopropanol. DNA-contaminating templates were removed by DNAse treatment using the DNA-free DNA Removal kit (Ambion). RNA concentrations were determined and equal amounts (0.9 μg) were used to generate cDNA using the TaqMan reverse transcription kit with random hexamers (Applied Biosystems), according to the manufacturer’s protocol. Equal amounts of cDNA (10 ng) were then analyzed by qPCR in triplicate using *Power* SYBR Green Master Mix (Applied Biosystems) and an Eppendorf Mastercycler RealPlex_2_ real-time PCR machine. Glyceraldehyde-3-phosphate dehydrogenase (*GAPDH)* RNA levels were quantified in parallel and used to normalize for the RNA isolation efficiency of each sample. Viral RNA copy number was calculated using a standard curve generated from HCMV BAC DNA, as above.

### Immunofluorescence assay (IFA)

MRC5 cells were infected with BAC-derived TB40/E lacking a fluorescent marker (38) at a multiplicity of 0.01 IU/cell. Cells that were harvested 6 dpi were washed three times with 37°C 1X PBS, and then fixed at 37°C with 2% paraformaldehyde for 15 min. The infected cells were washed with 1X PBS three times at room temperature (RT) and then permeabilized with 0.1% Triton X-100 for 15 min. Next, cells were washed with 0.2% Tween 20 in 1X PBS three times, blocked in 2% BSA with 0.2% Tween 20 in 1X PBS at RT for 1 h, and were then stained with the primary antibodies (pp71; 1:50 and STING; 1:100) at RT for 1 h. Cells were washed (0.2% Tween 20 in 1X PBS), and then stained with secondary antibodies (anti-mouse Alexa Fluor 488; anti-rabbit Alexa Fluor 594) and 4’,6-diamidino-2phenylindole (DAPI; to visualize nuclei) at RT for 1 h in the dark. Images were collected on an Olympus 1X81 microscope.

## RESULTS

### Distinct HCMV proteins are S-nitrosylated during infection of fibroblasts

Infection with viruses induces significant changes within a cell’s proteome, including PTM alterations. To monitor the changes in host cell protein S-nitrosylation in response to infection, we infected human fibroblasts (MOI = 3.0) with a BAC-derived, clinical isolate of HCMV that expresses a viral late protein, pp28 (encoded by *UL99*), containing an in-frame fusion to eGFP, so as to monitor viral replication. We collected protein lysates at 96 hours post-infection (hpi) in non-reducing conditions to preserve the protein S-nitrosylation modification of cysteines, since this modification is highly liable under standard reducing conditions. We then biotinylated the protein S-nitrosylated cysteines using a biotin switch assay (39), thus allowing for enrichment and subsequent peptide identification by tandem mass spectrometry (MS/MS) analysis. MS/MS analysis of fragmented peptides not only allows us to identify peptide amino acid sequences, but also affords us the platform to characterize the specific cysteines that are increased in mass due to the conjugated biotin. Thus, we can confirm protein S-nitrosylated of specific cysteines. Using this methodology, we analyzed three independent, biological replicates each of mock and infected MRC5 cells and identified more than 2000 peptides, representing 954 host proteins differentially S-nitrosylated in response to HCMV infection. Importantly, we identified S-nitrosylation of 23 viral peptides originating from 13 different viral ORFs at 96 hpi (Table 1), in which we also confirmed the presence of a modified cysteine in each peptide (represented as a lower case “c” in the peptide sequence, Table 1).

**Table 1.**
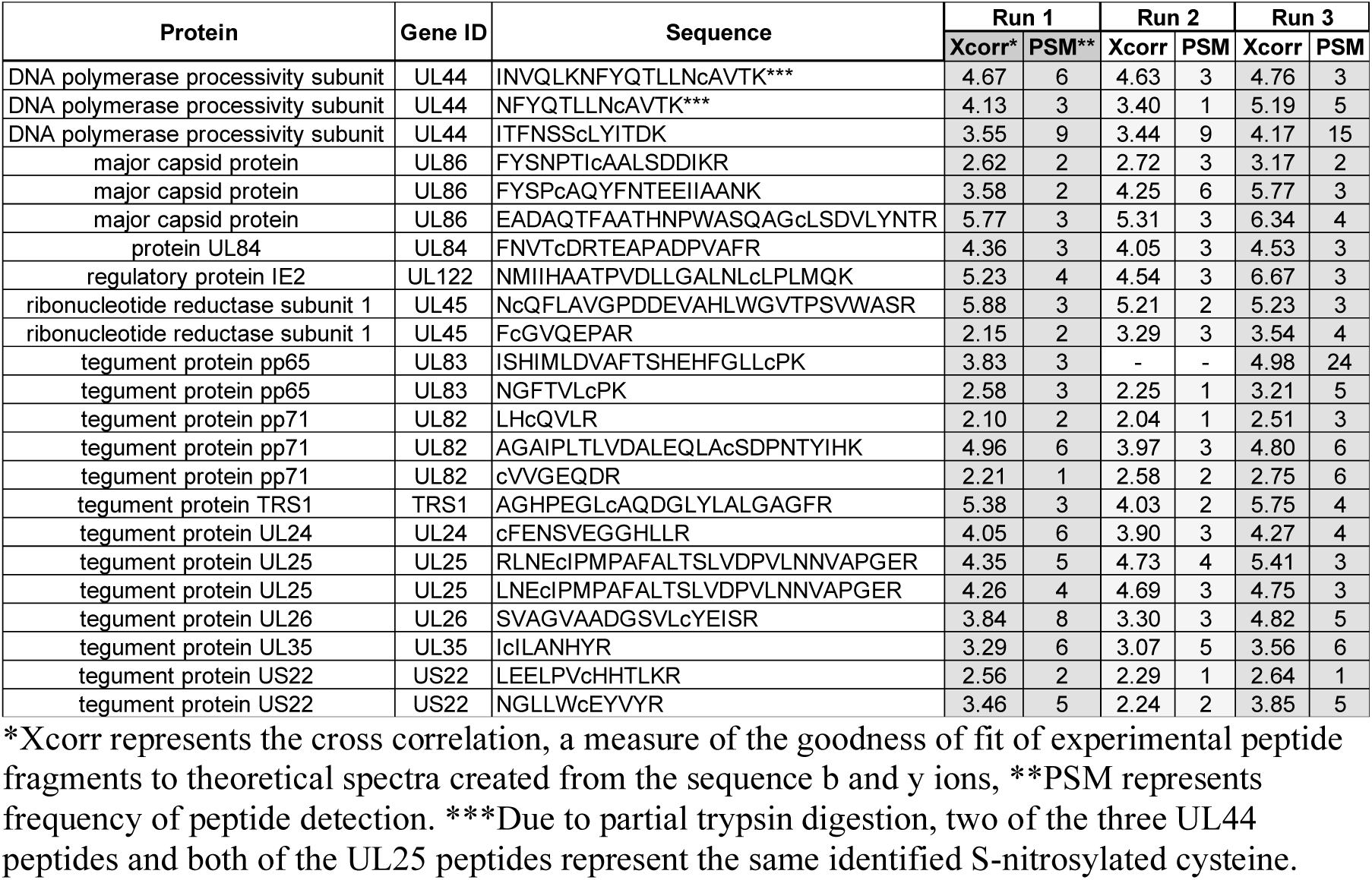
S-nitrosylated peptides of HCMV.

We were intrigued by S-nitrosylation of the *UL82*-encoded viral protein, pp71, which is important for mediating host antiviral responses involving the interferon pathway. In order to validate our MS/MS results, we repeated the infections and isolated equal amounts of whole cell lysates for biotin-switch labeling. We then enriched the S-nitrosylated, biotin-labeled proteins by coupling them to streptavidin beads and then eluted them for immunoblot blot analyses. A single protein band of the correct size and antigen specificity for HCMV pp71 was detected from the biotin-switch assay enriched proteins, thus confirming the modifications we identified in the MS/MS analyses (Fig 1A, top panel). In addition, we performed the reverse pull-down, in which lysates from the biotin switch reaction were immunoprecipitated with a pp71 specific antibody followed by blotting with an anti-biotin specific antibody. We identified biotinylated pp71 protein (Fig 1A, bottom panel), thereby verifying pp71 is specifically S-nitrosylated during infection of fibroblasts.

**Fig. 1.**
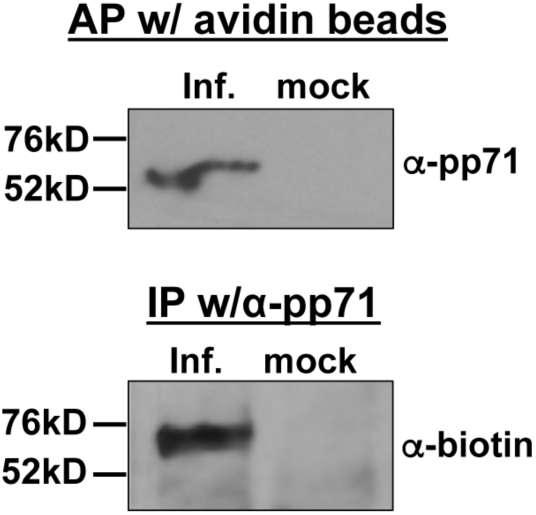
HCMV pp71 is protein S-nitrosylated during infection. NuFF-1 cells were infected at a multiplicity of 1.0 IU/cell with TB40/E-mCherry-UL99eGFP, and cells were harvested at 96 hpi. After the biotin-switch assay, lysates were affinity purified (AP) with avidin beads (top panel) or immunoprecipitated by pp71 specific antibody (bottom panel). Purified lysates were separated by 8% SDS-PAGE and transferred to a nitrocellulose membrane. Proteins were detected using a pp71 specific antibody (top panel) or antibody against biotin (bottom panel). n= 3; representative blots are shown.

### Mutation of protein S-nitrosylation identified cystine residues within pp71 does not inhibit viral production

To determine the biological function of the S-nitrosylated residue(s), we conducted mutational analysis of pp71. We introduced mutations within pp71 in the context of the viral BAC genome by changing each of the three identified S-nitrosylated cysteines to a structurally similar serine (C34S, C94S, and C218S). The resulting BAC was used to reconstitute the triple mutant pp71 virus (pp71-TM). To determine if the cysteine-to-serine (Cys-to-Ser) mutations within pp71 inhibit the viability of virus production, we monitored replication kinetics by viral growth analysis. To this end, we compared the single-step growth of the pp71-TM virus to that of the wild-type/parental virus in MRC5 fibroblasts, quantifying cell-free virus over a time course of 7 d (Fig 2A). We did not observe a significant viral growth defect when all three identified S-nitrosylated cysteines were mutated to serine within pp71 (pp71-TM). In fact, pp71-TM viruses exhibited a slight, yet consistent increase in both growth kinetics and cell-free viral yield when compared to wild-type virus (Fig 2A). To determine whether the triple mutation described above altered viral protein expression of pp71, we analyzed viral protein accumulation over a course of infection by immunoblot. We found that, pp71-TM infected fibroblasts displayed similar viral protein expression compared to its wild-type counterpart (Fig 2B). To determine when the Protein S-nitrosylation of cystines occurs on pp71, we analyzed the kinetics of this PTM on pp71 in relation to *de novo* pp71 protein synthesis. Fibroblasts were infected with wild-type HCMV and protein lysates were isolated at various timepoints. Lysates were either used for western blot analysis of total pp71 protein expression or used in a biotin switch assay, where S-nitrosylated proteins were captured with streptavidin beads and then analyzed by western blot to assess pp71 protein S-nitrosylation (Fig 3A). We observed similar kinetics between total pp71 expression and the appearance of the protein S-nitrosylation PTM on pp71 suggesting the modification on the protein occurs in a similar time frame as protein synthesis. Thus, we focused on characterizing the biological effects of pp71 S-nitrosylation.

**Fig. 2.**
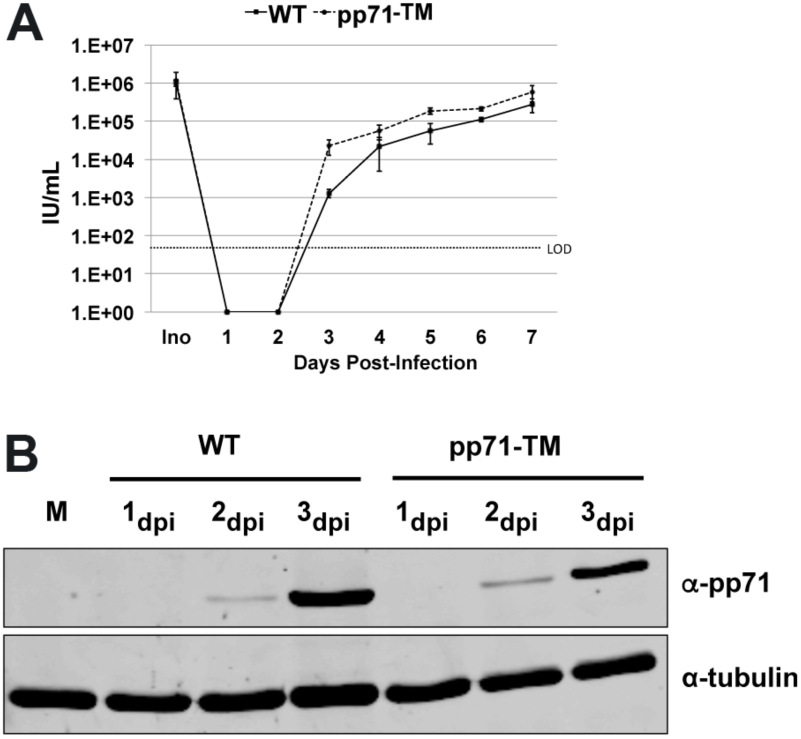
S-nitrosylation defective viruses have similar growth kinetics compared to WT virus. (**A**) NuFF-1 cells were infected at a multiplicity of 1 IU/cell with TB40/E-mCherry-UL99eGFP virus (WT), or a recombinant virus containing three point mutations in pp71 ORF at amino acids C34S, C94S, and C218S (pp71-TM). Media containing cell-free virus was collected over the indicated time course, and virus yields were determined by TCID_50_ assay on NuFF-1 cells. Ino = inoculum; LOD = Limit of detection. Samples were analyzed in triplicate. Error bars represent the standard deviation of the replicates. (**B**) Total, *de novo* expression of wild-type pp71 or pp71-TM was monitored by immunoblot. NuFF-1 cells were infected at a multiplicity of 1 IU/cell with WT or pp71-TM. Cells were harvested at the indicated time points after infection and the proteins separated by 8% SDS-PAGE and transferred to a nitrocellulose membrane. Cellular tubulin levels served as a control for equal protein loading. n= 3, representative blots are shown.

**Fig. 3.**
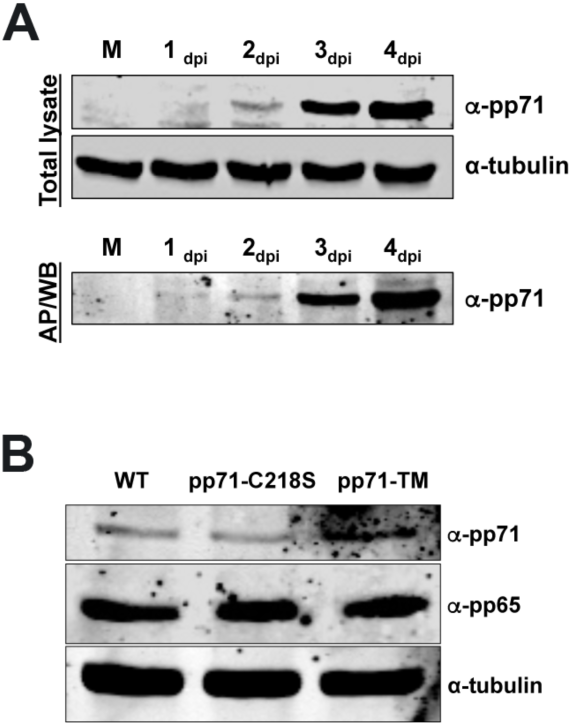
Protein S-nitrosylation defective pp71 is expressed with wild-type kinetics and incorporated into tegument. (**A**) NUFF-1 cells were infected at a multiplicity of 1 IU/cell with WT virus. Cells were harvested at the indicated time points after infection and the protein lysates were either separated by SDS-PAGE gel electrophoresis and blotted to nitrocellulose membrane (top panel; Total lysate) or used in a biotin switch assay, affinity purified on streptavidin beads (AP) and then separated by SDS-PAGE and transferred to a nitrocellulose membrane (bottom panel; AP/WB). Cellular tubulin levels serve as a control for equal protein loading. N = 3; representative blots are shown. (**B**) MRC5 cells were infected at a multiplicity of 30 IU/cell with the TB40/E-mCherry-UL99eGFP virus, pp71-C218S, or pp71-TM. Cells were harvested 6 hpi to detect pp71 and pp65 delivery into newly infected cells. Total cell lysates were separated by 8% SDS-PAGE and analyzed by western blot. Cellular tubulin levels served as a control for equal protein loading. n= 3; representative blots are shown.

pp71 is a multifunctional protein critical for efficient viral infection. Among its many roles, pp71 interacts with pRb resulting in its degradation, mediated by the consensus pRb binding motif LxCxD in pp71, where the central cysteine is located at amino acid 218 (14). Our proteomic analysis revealed pp71 cysteine 218 (C218), the central cysteine of the pRb binding motif, is S-nitrosylated (Table 1). To elucidate the biological consequence(s) of C218 S-nitrosylation, we introduced a single Cys-to-Ser mutation of C218, C218S, into the wild-type BAC genome. We first determined whether C218 S-nitrosylation was critical for tegument delivery of pp71 upon infection. We generated recombinant viruses and assessed their ability to deliver tegument pp71 as well as the well characterized tegument protein, pp65 as a control. At high moi infection of fibroblasts (MOI = 30), we detected wild-type, as well as pp71-TM and pp71C218S at 6 hpi (Fig 3B). Although pp71 is a late kinetic viral protein, its detection in newly infected cells early post-infection (Fig 2B) and prior to *de novo* synthesis is consistent with tegument-delivered pp71 (40). Together, these results suggest the ability to inhibit protein S-nitrosylation of pp71 did not significantly alter the growth kinetics or production of viral progeny when compared to the parental virus. In addition, mutation of the identified protein S-nitrosylation modified cysteines did not alter the expression, tegument loading, or delivery of pp71 to newly infected fibroblasts.

### Protein S-nitrosylated pp71 interacts with STING

We next assessed the importance of protein S-nitrosylation for known biological functions of pp71. Recent studies found pp71 could inhibit STING activation (21). Inhibition of STING may be a universal feature of successful viruses, as both adenovirus E1a and human papilloma virus E7 also inhibit STING anti-viral functions (41). E1a and E7 mediate these interactions through their conserved LxCxE pRb binding domain. As our analysis identified the central cysteine in the pp71 LxCxD domain was S-nitrosylated during infection, we next asked whether this PTM affected pp71:STING interactions. HEK293T cells lacking endogenous STING (42) were co-transfected with either empty vector or a vector expressing HA-tagged STING (STING-HA) (34) in conjunction with either FLAG-tagged wild-type pp71 (FLAG-WT-pp71) or FLAG-tagged pp71 bearing a C218S mutation within the pRb binding domain (FLAG-pp71-C218S). First, we confirmed the expression of each epitope-tagged protein by immunoblot analysis (Fig 4A). Transiently co-transfected cell lysates precipitated with anti-FLAG antibody followed by immunoblot revealed both FLAG-WT-pp71 and FLAG-pp71-C218S each interact with STING-HA (Fig 4A, top panel). We also found the same interaction between FLAG-pp71-C218S or FLAG-WT-pp71 with STING-HA when we reversed the order of immunoprecipitation, using an HA antibody followed by immunoblot with an anti-FLAG antibody (Fig 4A, bottom panel). We observed STING-HA immunoprecipitated FLAG-pp71-C218S with greater efficiency than FLAG-WT-pp71, suggesting an increased binding affinity of protein S-nitrosylation-defective pp71 for STING-HA (Fig 4A, bottom panel). Collectively, these results suggest STING interacts with pp71 and the interaction may occur with greater affinity when pp71 C218 within its pRb-binding domain cannot undergo protein S-nitrosylation.

**Fig. 4.**
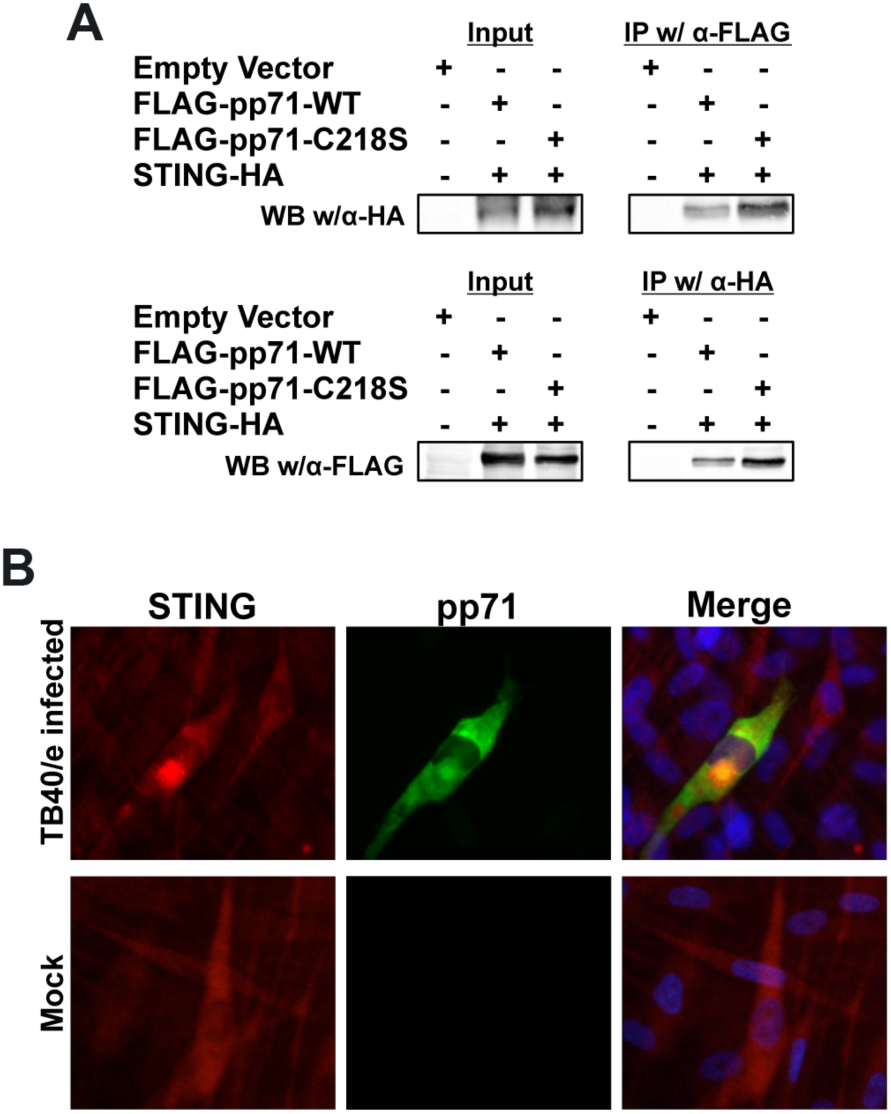
pp71 interacts with STING *in vitro* and co-localizes with STING during infection. (**A**) HEK-293 cells were co-transfected with pCDNA-STING-HA (STING-HA) and pCMSeGFP-flag-pp71-WT (FLAG-pp71-WT) or pCMSeGFP-flag-pp71-C218S (FLAG-pp71-C218S). After 72 h, total cell lysates were collected. Equal amounts of total cell lysates were incubated with α-HA or α-FLAG affinity beads. The bound complexes were washed, eluted, separated by 8% SDS-PAGE and immunoblotted (IB) with an α-FLAG or an α-HA antibody. n=3; representative blots are shown. (**B**) MRC5 cells were infected at a multiplicity of 0.01 IU/cell with the HCMV TB40/E-mCherry-UL99-eGFP virus (upper panels) or mock infected (lower panels). At 6 dpi, cells were fixed with 2% PFA, and were stained with antibodies specific for STING (rabbit polyclonal) and pp71 (mouse monoclonal). STING-HA was detected using an Alexa-594 conjugated antibody that recognizes the Fc domain of rabbit polyclonal antibodies (red) and pp71 was detected using an Alexa-488 conjugated antibody that recognizes the Fc domain of mouse monoclonal antibodies (green). DAPI (blue) was used to visualize the nuclei. n=3; representative images shown.

To test our hypothesis that pp71 interacts with STING in the context of infection, we performed immunofluorescent assays (IFAs) to monitor co-localization of pp71 and endogenously expressed STING. Human fibroblasts were either mock-infected or infected with a BAC-derived parental, TB40/e clone 4 (38) lacking a fluorescent gene. At 6 dpi, cells were fixed and stained with antibodies that recognize pp71 and cellular STING. STING was observed in both mock-infected and infected cells, whereas pp71 was expressed only in infected cells (Fig 4B). Intracellular STING localization differed in infected cells compared to mock-infected cells. In infected fibroblast, STING localized to a perinuclear region where pp71 was also enriched, suggesting that the two proteins may co-localize during infection (Fig 4B), within subcellular regions consistent with the localization of the viral assembly compartment (43).

Enhanced STING staining within infected cells leads one to question whether viral infection results in the re-localization of endogenous STING or increased STING expression in infected host cells. To test this, we infected NuFF-1 cells with either WT HCMV or recombinant virus expressing pp71-C218S (MOI = 1.0). We collected total cell lysates at time points relevant for tegument-delivered pp71 (Fig 3B), as well as at time points consistent with *de novo* protein synthesis of pp71 (Fig 3A). Neither tegument-delivered nor *de novo* protein synthesis of WT or mutant pp71 increased the relative expression of endogenous STING compared to that of α-tubulin (Figs 5A & B). These results reveal pp71 does not regulate endogenous STING expression levels, suggesting the increased STING staining in the IFA was most likely due to altered subcellular localization of STING.

**Fig. 5.**
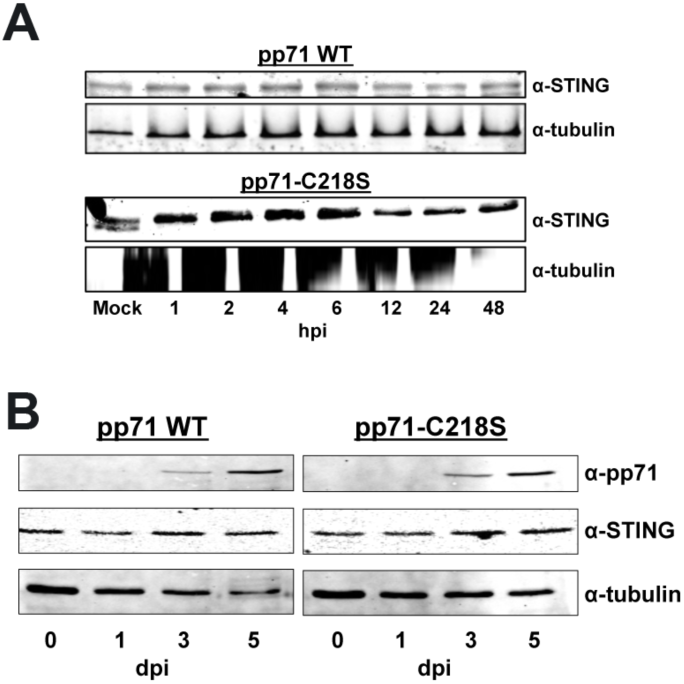
pp71 does not alter the protein expression levels of STING at early or late times during lytic infection. MRC5 cells were infected at a multiplicity of 1 IU/cell with WT or pp71-C218S. Infected cells were harvested over (**A**) 48 h prior to *de novo* pp71 expression or (**B**) 5 d, when *de novo* pp71 is expressed. (**A, B**) Total protein (20µg) was separated by SDS-PAGE and transferred to a nitrocellulose membrane. Cellular tubulin is shown as a loading control. n=3; representative blots shown.

### pp71 protein S-nitrosylation limits its ability to undermine the STING antiviral pathway

Since we observed small but reproducible increases in both growth kinetics and absolute titer following lytic infection with the S-nitrosylation-deficient recombinant virus, pp71-C218S, when compared to WT virus, we next investigated the biological impact of pp71 S-nitrosylation on the STING pathway in the context of viral infection. To this end, we generated a repair of the TB40/e-mCherry-pp71-C218S mutant, in which we restored the serine mutation to cysteine, thus reverting the sequence to wild-type (C218S*rev).* The use of this revertant ensures any phenotype we observed with the pp71p-C218S recombinant virus was not caused by off-site mutations due to the recombineering. To determine the effect of STING activation on viral replication, we treated MRC5 cells with 2’3’cGAMP and then infected the cells with WT, pp71-C218S, or pp71-C218S*rev*. 2’3’cGAMP is a natural ligand for STING activation (44), and a dose response curve showed treatment of MRC5 fibroblasts with 10µM 2’3’cGAMP was sufficient to activate STING signaling (Supplementary Fig 1A) and had no adverse effect on cell viability, as determined by MTT assay (Supplementary Fig 1B). We then monitored cell-associated HCMV genomes in MRC5s over a time course of viral infection. As expected, activation of the potent antiviral STING pathway yielded low copies of cell-associated viral genomes during WT or pp71-C218S*rev* infection (Fig 6A). However, infection with the S-nitrosylation-deficient pp71 mutant virus, pp71-C218S, resulted in a significant increase in cell-associated viral genomes (Fig 6A), suggesting S-nitrosylation prevents pp71-mediated inactivation of STING.

**Fig. 6.**
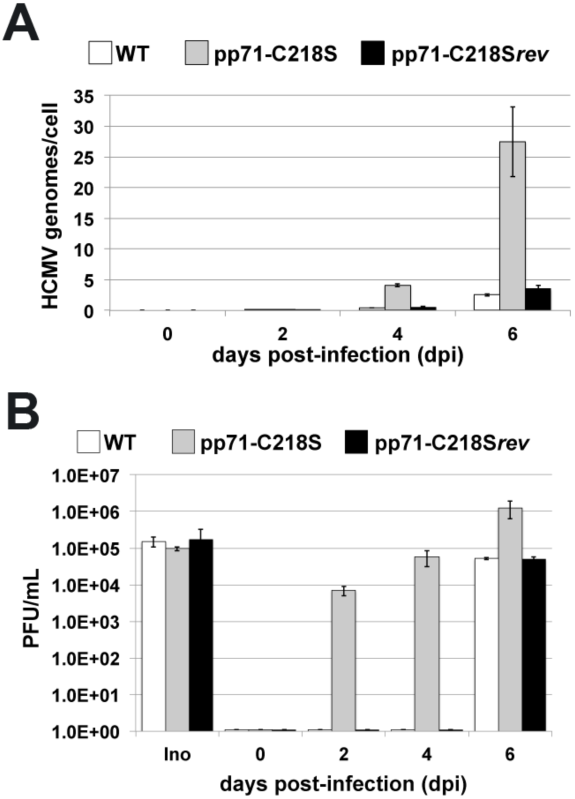
Activation of the STING pathway inhibits DNA replication and viral production of pp71-WT and pp71-C218S*rev* but not pp71-C218S. (**A**) MRC5 cells were infected with either WT, pp71-C218S, or pp71-C218S*rev* (MOI=1) in the presence of 10uM 2’3’cGAMP. DNA was isolated over the indicated days, and genome copy number was measured by qPCR using eGFP-specific primers to detect viral genomes and normalized to human mdm2; (**B**) Parallel cultures were treated as in (**A**) and media was collected over the indicated days. Cell-free virus accumulation was quantified by TCID_50_ analysis. Ino=inoculum. (n = 3).

We extended this analysis to monitor the impact of pp71 protein S-nitrosylation on accumulation of infectious, cell-free virus in the presence of activated STING by single-step growth analysis. MRC5 cells were infected with WT, pp71-C218S, or pp71-C218S*rev* (MOI = 1) in the presence of 2’3’cGAMP. Cell-free viral supernatants were collected over time for quantification of viral production by TCID_50_. Analysis of the initial infectious inoculum confirmed an equal amount of each virus was used in the experiment (Fig 6B, Ino). We observed markedly diminished viral yield in both the WT- and pp71-C218S*rev*-infected fibroblasts in the presence of 10µM 2’3’cGAMP up to day 4 of the experiment (Fig 6B). However, by 6 dpi both the WT- and pp71-C218S*rev*-infected cells began to produce infectious progeny from the 2’3’cGAMP treated cells (Fig 6B). Importantly, infection with pp71-C218S, the protein S-nitrosylation-deficient mutant virus, efficiently accumulated infectious progeny even in the presence of 2’3’cGAMP across the time course of infection. Together, these results suggest protein S-nitrosylation-deficient pp71 at position C218 undermines potent STING activation compared to wild-type pp71.

### Activated STING inhibits viral transcription of WT virus but not of pp71-C218S

To determine the stage of viral replication affected by pp71 S-nitrosylation following STING activation, we monitored the accumulation of a subset of HCMV transcripts within MRC5 cells. To this end, these fibroblasts were infected with WT, pp71-C218S, or pp71-C218S*rev* in the presence or absence of STING activation. Viral transcripts *UL122*, *UL54*, and *UL32* were selected as representative genes of each infection stage (e.g.: immediate early (IE), early (E), and late (L), respectively (45)). In the absence of STING activation, the expression and accumulation of HCMV transcripts were similar to each other when compared across each virus infection (Fig 7A, upper panel). As expected, activation of the STING pathway with 2’3’cGAMP treatment prior to infection significantly inhibited the expression and accumulation of HCMV transcripts in the fibroblasts that were infected with either WT or pp71-C218S*rev* viruses. However, 2’3’cGAMP treatment did not impair the transcription of these viral genes in the pp71-C218S infected fibroblasts (Fig 7A, lower panel). Indeed, the viral transcript levels in the pp71-C218S infected cells in the presence of 2’3’cGAMP (Fig 7A, lower panel) mirrored the levels we observed during HCMV infection in the absence of activated STING (Fig 7A, upper panel). These data indicate STING activation inhibits HCMV infection and expression of S-nitrosylation-deficient pp71 effectively undermines this anti-viral response.

**Fig. 7.**
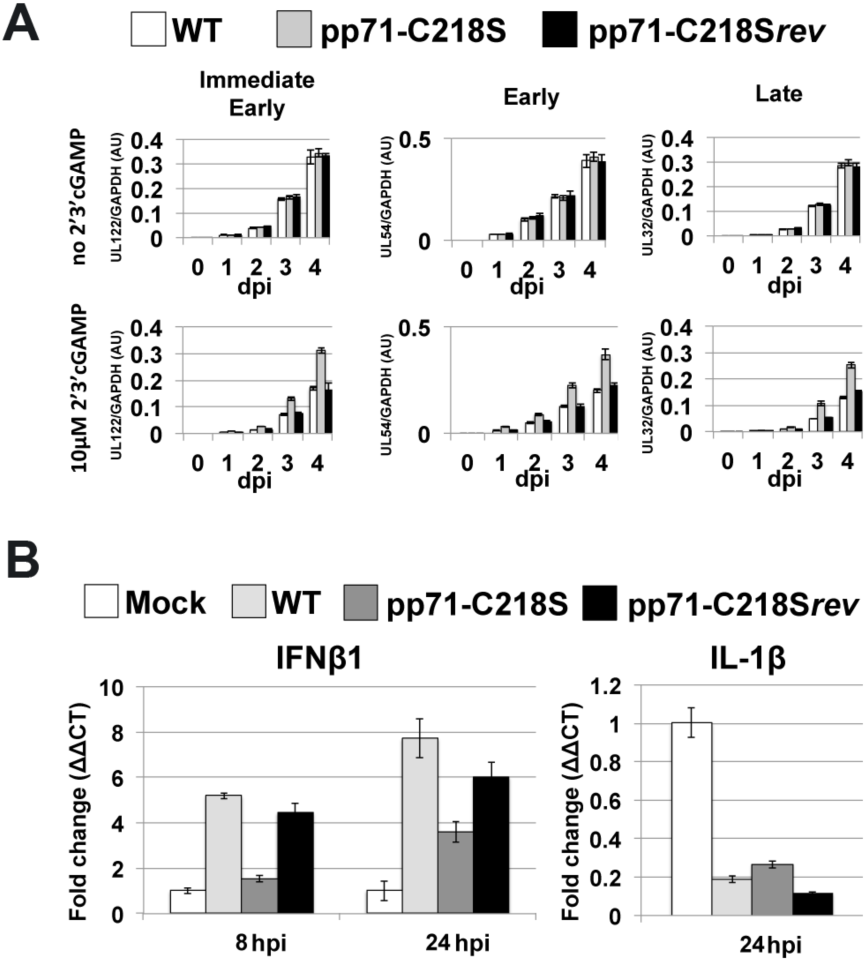
STING activation impacts viral RNA accumulation in WT- and pp71-C218S*rev*-infected cells, but not pp71-C218S-infected cultures. (**A**) MRC5 cells infected with either WT, pp71-C218S, or pp71-C218S*rev* (MOI=1) in the absence or presence of 10uM 2’3’cGAMP (added at time of infection). Total RNA was isolated from the cultures over the indicated times. Transcripts IE (*UL122*), E (*UL54*), and L (*UL32*) were profiled by RT-qPCR using transcript-specific primers and normalized to cellular *GAPDH*. Results are shown as mean fold-change in the mRNA levels for each of three different kinetic classes of viral transcripts within cells infected by the listed viruses in the presence of vehicle (top) or 10µM 2’3’cGAMP (bottom). AU, arbitrary units. (**B**) MRC-5 cells were mock-infected or infected with WT, pp71-C218S, or pp71-C218S*rev* (MOI=1) in the presence of 10µM 2’3’-cGAMP. Total RNA was isolated at 8 or 24 hpi. *IFNβ1* and *IL-1β* transcripts were profiled by RT-qPCR using transcript-specific primers and plotted as ΔΔCT relative to the control transcript, *GAPDH*. Results are shown as mean fold-change in the mRNA levels of *IFNβ1* and *IL-1β* in the cells.

### Protein S-nitrosylation of pp71 impacts STING induced transcriptional activation

Activated STING promotes a potent antiviral state within infected cells, leading to the induction of both nuclear factor kappa-light-chain-enhancer of activated B cells (NFκB)- and interferon regulatory factor 3 (IRF3)-mediated transcription (23). This leads to accumulation of cytokines and interferon genes, both of which limit viral replication. To determine whether protein S-nitrosylation of pp71 impacts these transcriptional pathways after STING activation, 2’3’cGAMP treated MRC5 cells were infected with WT, pp71-C218S, or pp71-C218S*rev* and the expression of NFκB- and IRF3-responsive genes was quantified by RT-qPCR. As expected, expression of IFNβ1 was upregulated at both 8 and 24 hpi in WT and pp71-C218S*rev* infected cells following STING activation, however, induction of IFNβ1 was suppressed in pp71-C218S infected cells (Fig. 7B). Expression of the NFκB-responsive IL-1β transcript was equally suppressed with each of the viruses used, suggesting the inhibition was independent of C218 S-nitrosylation. Previous studies showed HCMV-encoded proteins, pUL26 and pp65 inhibit the NFκB pathway (46, 47). As pUL26 and pp65 were expressed in the three viruses used in this stud, we desired to evaluate pp71 protein S-nitrosylation in the absence of other viral proteins. However, our observations strongly suggest the induction of IFNβ1 transcription following the activation of STING is inhibited by pp71 deficient for protein S-nitrosylation at C218.

### pp71 functions to inhibit STING-mediated transcription independent of other viral factors

We have shown blocking protein S-nitrosylation of pp71 is necessary for efficient inhibition of STING-induced transcription. Thus, we next determined if blocking this PTM is sufficient by assessing pp71 in the absence of viral infection. To this end, MRC5s stably transduced with lentiviral constructs that allow for the stable expression of either WT-pp71 (M5-pp71_WT_) or S-nitrosylation-deficient pp71, pp71-C218S (M5-pp71_c218S_) were generated (Fig 8A). We next determined whether exogenously expressed pp71 was S-nitrosylated in the absence of HCMV infection. We performed the biotin-switch assay on lysates from M5-pp71_WT_ transduced cells and found pp71 was indeed S-nitrosylated (Fig 8B), suggesting this PTM occurs independent of any additional viral proteins. We did not perform this analysis on the M5-pp71_c218S_ cells as this mutant pp71 construct still contains the remaining two cysteines (C34 and C94) that we showed are nitrosylated (Table 1). Thus, this mutant would still display protein S-nitrosylation, as this methodology does not distinguish the location of the PTM.

**Fig. 8.**
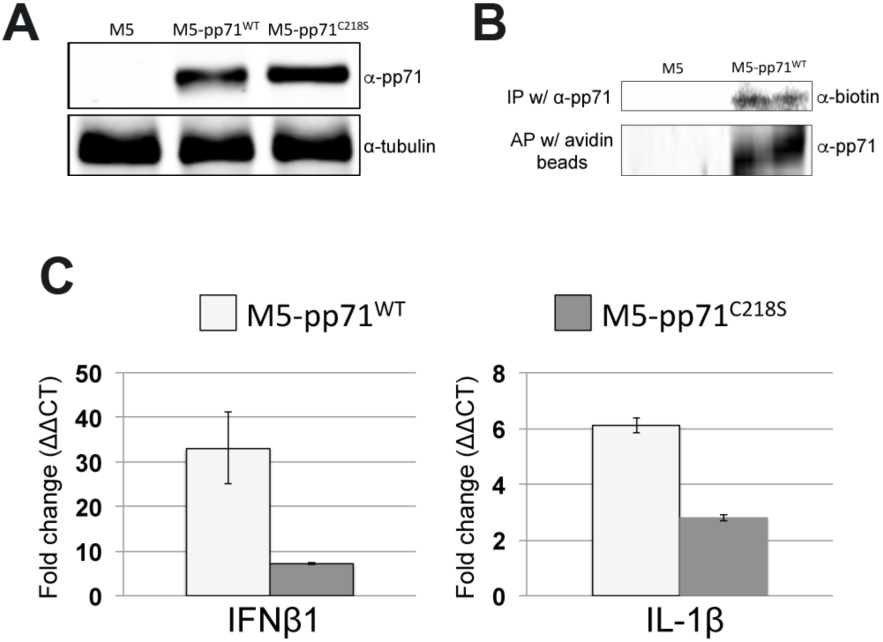
S-nitrosylation of pp71 is independent of viral infection and impacts STING-induced transcription in the absence of additional HCMV factors. (**A**) Recombinant retroviruses, pLXSN-pp71-WT or pLXSN-pp71-C218S were used to transduce MRC5 cells that were then selected in G418 for two cell doublings yielding the parental MRC5 cells (M5), M5-pp71_WT_, and M5-pp71_C218S_. Cell lysates were isolated and total protein (20µg) was separated by SDS-PAGE and then transferred to a nitrocellulose membrane. Immunoblotting was performed with an α-pp71 and α-tubulin, which served as a loading control; representative blots are shown. (**B**) Lysates from M5 and M5-pp71_WT_ were harvested and used in a biotin switch assay. Lysates were either affinity purified (AP) with avidin beads or immunoprecipitated (IP) with pp71 antibody. Proteins were separated by SDS-PAGE, then transferred to nitrocellulose membrane. Protein S-nitrosylated pp71 was detected by α-biotin or α-pp71 antibody. n=3; representative blots are shown. (**C**) 2’3’-cGAMP (10µM) was added to M5-pp71_WT_ and M5-pp71_C218S_, and 24 h later, total RNA was isolated. *IFNβ1* and *IL-1β* transcripts were profiled by RT-qPCR using transcript-specific primers and plotted as ΔΔCT relative to the control transcript, *GAPDH*. Results are shown as mean fold-change in the mRNA levels of *IFNβ1* and *IL-1β* in M5-pp71_WT_ and M5-pp71_C218S_ cells.

To determine whether stable expression of either WT or mutant pp71 impacts interferon induction in fibroblasts in the absence of infection, we treated the transduced cells with 2’3’cGAMP and monitored transcription of STING-responsive transcripts by RT-qPCR. We found the M5-pp71_c218S_ cells displayed a significant reduction of IFNβ1 when compared to the M5-pp71_WT_ cells (Fig. 8C). As we performed these experiments in the absence of any other HCMV proteins that may impact NFkB, we were able to characterize the impact of pp71 S-nitrosylation on the NFkB-mediated pathway by quantifying IL-1β express. In support of our hypothesis that pp71 S-nitrosylation affects the inhibition of the STING pathway, our data revealed IL-1β expression was significantly inhibited by overexpression of pp71-C218S compared to WT-pp71 (Fig. 8C). Together, these results reveal pp71 inhibited STING-mediated signal transduction, as previously reported, thereby attenuating the STING antiviral response. Importantly, pp71 S-nitrosylation impeded this STING inhibition, and a single amino acid substitution within the pRb-binding domain of pp71 was sufficient to relieve this repression.

## DISCUSSION

We undertook an unbiased approach to identify and characterize the HCMV viral proteome that undergoes protein S-nitrosylation during lytic infection. As this modification is reversible, we hypothesized this PTM would play distinct roles at different points of the lytic life cycle. To this end, we characterized the S-nitrosylated proteome at a time point during infection in which viral DNA has initiated and assembly of infectious progeny has begun (96 hpi). We chose this time point for two reasons: 1) most viral proteins are expressed at high levels at this time, and 2) protein S-nitrosylation is a known host response to stress (48). It should be noted this methodology may overlook viral proteins with short half-lives, transient protein S-nitrosylation modifications, and peptides from viral ORFs not present at this time during infection. In three biological replicates, we consistently identified 13 viral proteins from each kinetic class of HCMV proteins that are S-nitrosylated (Table 1) with more than half of these peptides originating from tegument proteins.

Of particular interest was the identification of three distinct S-nitrosylation sites within pp71. The tegument viral protein pp71 has multiple, critical biological functions which promote efficient viral replication by undermining host antiviral responses. Specifically, tegument delivered pp71 functions to degrade hDAXX as well as pRb, thus relieving repression of viral transcription and promoting entry into the cell cycle of the infected host cell, respectively (14–17, 49). Thus, evaluating the impact of pp71 protein S-nitrosylation was a primary focus. With advancements in bacterial artificial chromosome (BAC) recombineering technology, we are able to mutate and evaluate the requirement of specific protein motifs in viral proteins within the context of an active infection. We mutated each of the S-nitrosylated cysteines we identified in pp71 to the closely related amino acid serine, which is incapable of protein S-nitrosylation. Unexpectedly, mutation of the modified cysteines within pp71 resulted in a viable virus that replicated with a modest, yet consistently better efficiency than wild-type virus (Fig 2A). We had hypothesized any deleterious mutation would result in a growth defect in the virus, especially as one of the mutated cysteines was located within the LxCxE/D domain previously identified in pp71 to mediate the pRb binding functions (14). However, mutations of all three identified protein S-nitrosylated cysteines to serines within pp71 resulted in nearly equal amounts of viral protein accumulation when compared to wild-type (Fig 3A) and no apparent defect in tegument loading of pp71 into cell-free virions (Fig 3B) suggesting that these mutations resulted in no apparent biological defect within highly permissive fibroblasts.

More recently, pp71 expression has been implicated in undermining the STING-induced response in HCMV-infected cells (21). STING is an adaptor protein involved in mediating the response between activated cytoplasmic DNA sensors and the type I interferon response pathway. STING activation, induced by encountering the secondary signaling mediator 2’3’cGAMP, promotes a potent antiviral state resulting in NFkB-mediated transcriptional activation of cytokines and IRF3-mediated transcriptional activation of interferon genes. Thus, pp71’s ability to undermine STING activation is critical for efficient viral replication. Importantly, proteins encoded by other viruses undermine STING responses. Human papilloma virus E7 and human adenovirus E1a both interact with STING, and these interactions require the LxCxE/D pRb-binding motif found in each of these viral proteins (41). Importantly the best-characterized protein S-nitrosylation motif is the I/LxCx_2_E/D motif, initially characterized in regulating the function of glyceraldehyde-3-phosphate dehydrogenase (GAPDH) within the interferon-activated inhibitor of translation (GAIT) complex (50). Protein S-nitrosylation regulates the function of the GAIT complex, thus restoring translation of specific transcripts during times of interferon activation (51). This motif differs from the canonical pRb binding motif by only the spacing of a single amino acid. Thus, we hypothesized the protein S-nitrosylation of the central cysteine within pp71’s pRB-binding domain is critical for the interaction of this viral protein with STING.

We further investigated the biological impact of pp71 S-nitrosylation on the STING pathway. We observed, as previously reported (21), pp71 immunoprecipitated with STING, but we noted increased precipitation of the two proteins when we mutated the central cysteine in pp71’s pRb-binding domain to serine, thus blocking S-nitrosylation (Fig 4A). This suggests that S-nitrosylation of pp71 may be inhibitory to its biological functions. The interaction of pp71 with STING did not result in degradation of this host protein (Fig 5A&B), suggesting the mechanism of inhibiting the interferon pathway differs from how pp71 inhibits the transcriptional inhibitor, hDAXX.

To evaluate the impact of pp71 S-nitrosylation on the STING pathway, we assessed viral replication in the presence of the natural ligand for STING activation, 2’3’cGAMP. In support of our model, we observed inhibition of wild-type HCMV during activation of the STING pathway, but this attenuation was less pronounced when pp71 S-nitrosylation was blocked by the Cys-to-Ser mutation (Fig 6A&B). The net effect was a decrease in the production of antiviral cytokine and interferon transcripts (Fig 8). This led us to the novel interpretation that host-directed S-nitrosylation of virulence factors may function as an additional antiviral measure, thus limiting viral replication. As viral proteins essential for replication of other unrelated viruses (e.g. human papillomavirus and human adenovirus) have conserved motifs to pp71’s protein S-nitrosylation site through which they interact with STING, we favor a model where host cell mediated PTMs function to limit additional pathogens beyond HCMV, thus suggesting a more universal function for protein S-nitrosylation.

Why would HCMV pp71 evolve to maintain a motif that limits its capacity to replicate in the face of potent antiviral responses? Data suggests S-nitrosylation of cysteines within proteins is not random and that this modification is mediated by specific nitrosyltransferases selective for specific, conserved cysteines (52). One could speculate that there would be a benefit to pp71 altering this central cysteine, however, the pp71’s pRb-binding function is important during infection of certain cell types (53), thus maintaining this cysteine has a fitness benefit for the virus. It will be interesting to discern the extent of protein S-nitrosylation of different viral proteins to determine if this mechanism of host-mediated antiviral responses is more universal and potentially exploitable as an antiviral therapy. In sum, we have identified and characterized a novel pp71 PTM that functions as an additional host-mediated antiviral measure in response to infection. The balance between a successful infection and an abortive infection pivots on the ability of a virus to undermine host countermeasures to infection. We favor a model in which host cells are able to sense the initial shift towards successful viral replication and in response, the host cell induces pathways resulting in the accumulation of post translational modifications that incapacitate or limit potent viral virulence factors, such as pp71. In doing so, host cells can slow viral spread. Identifying the direct factors that dictate these responses will be of high value.

## ACKNOWLEDGEMENTS

The authors would like to thank Belinda Willard of the Cleveland Clinic Proteomics and Metabolomics Core Facility for her assistance with the proteomics reported in this manuscript.

## SUPPLEMENTL FIGURE LEGENDS

**Sup Fig 1.**
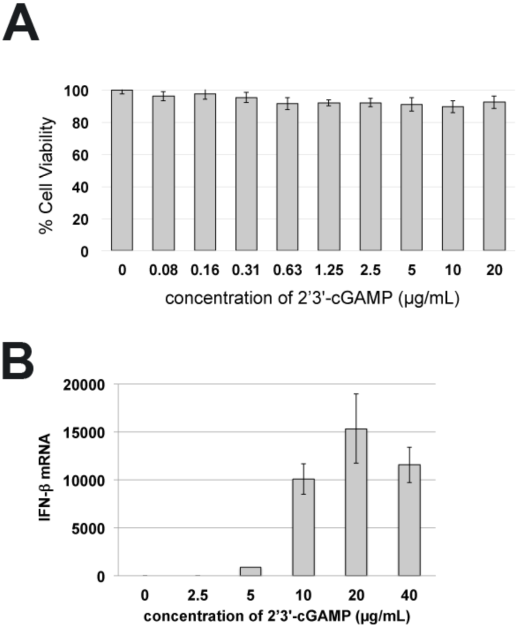
Fibroblasts treated with *2’3’*-cGAMP are viable and express IFNβ1. (A) MRC5 cells were treated with indicated concentrations of 2’3’-cGAMP, 48 hrs later cell viability was assessed by quantifying the metabolic activity of the treated cells using a NADH-dependent cellular oxidoreductase enzyme assay (MTT assay). MTT conversion results were normalized to the vehicle treated conditions and graphed results are shown as percent cell viability. Results are shown as the mean of three independent biological replicates. (B) MRC5 cells were treated with indicated concentrations of 2’3’-cGAMP, and 24 h later, total RNA was isolated. *IFNβ1* transcripts were profiled by RT-qPCR using transcript-specific primers and plotted as ΔΔCT relative to the control transcript, *GAPDH*. Results are shown as mean fold-change in the mRNA levels.

**Table S1.**
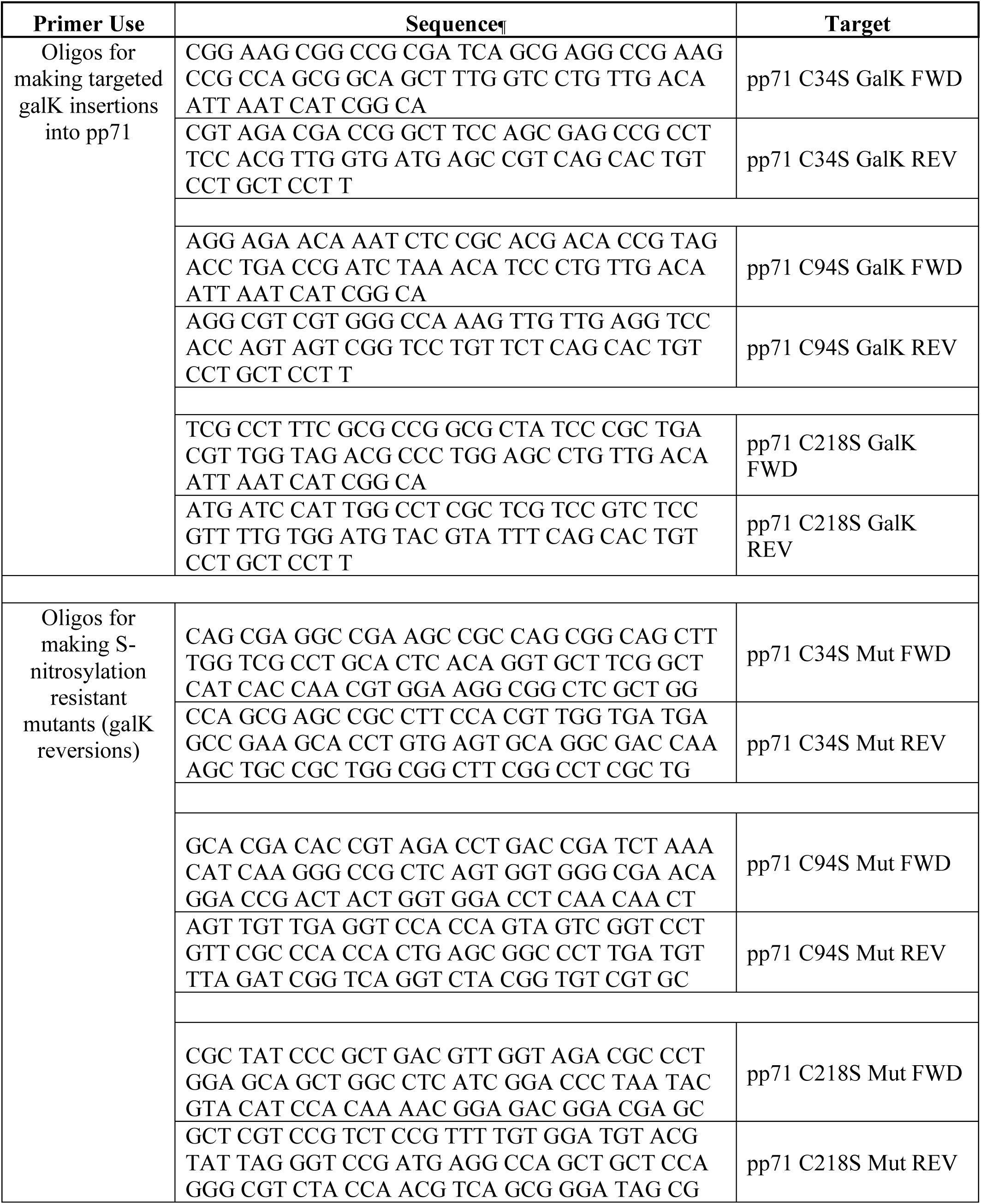
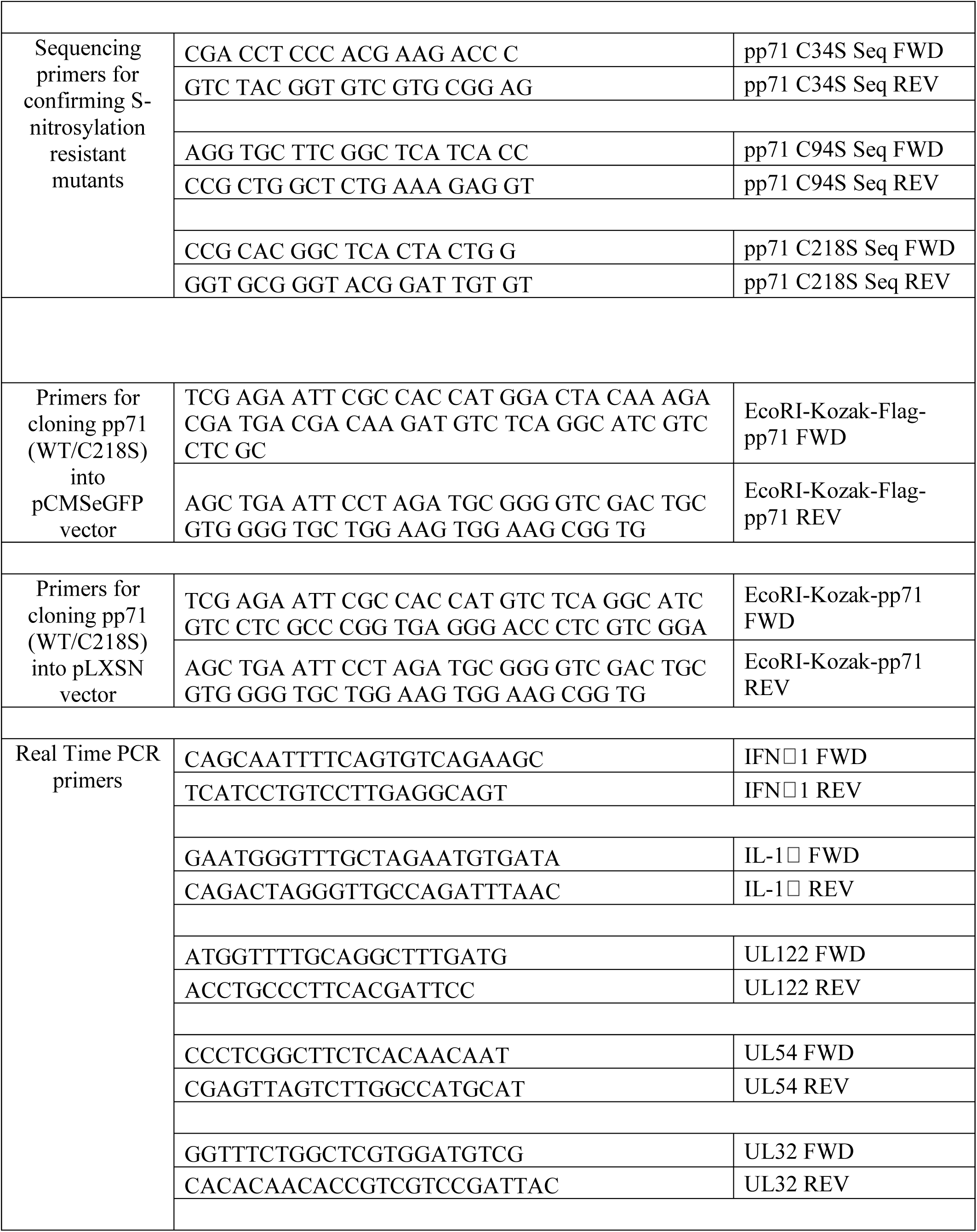
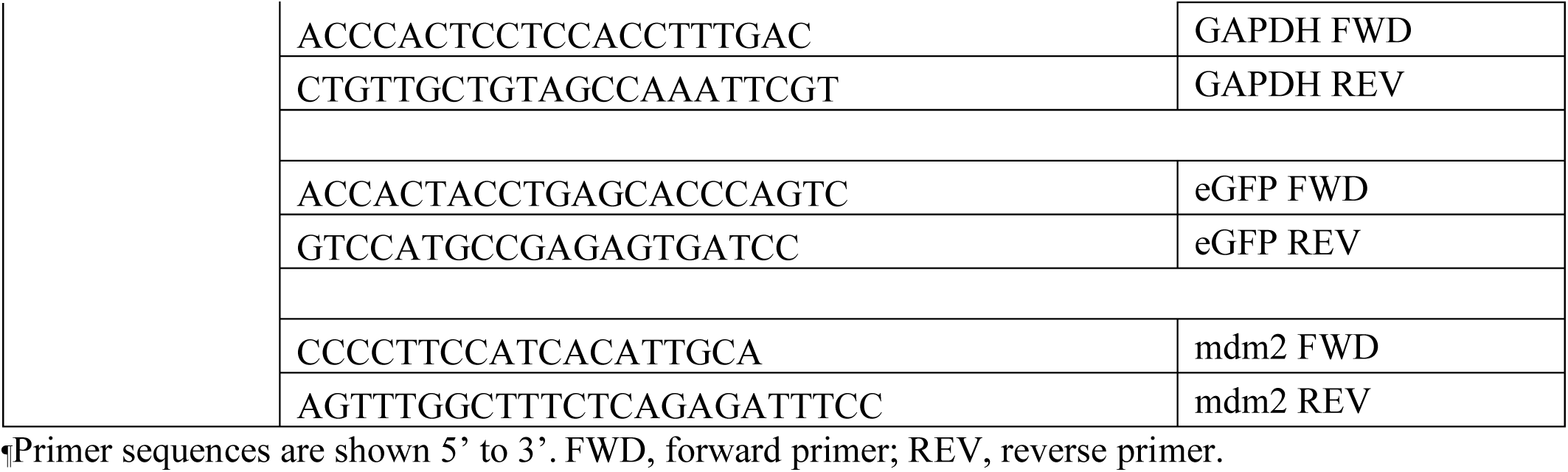
Oligonucleotides used in this study.

## REFERENCES

1. Britt WJ, Prichard MN. 2018. New therapies for human cytomegalovirus infections. Antiviral Res 159:153–174.

2. Britt W. 2008. Manifestations of human cytomegalovirus infection: proposed mechanisms of acute and chronic disease. Curr Top Microbiol Immunol 325:417–470.

3. McCormick AL, Mocarski Jr ES. 2007. Viral modulation of the host response to infection. In Arvin A, Campadelli-Fiume G, Mocarski E, Moore PS, Roizman B, Whitley R, Yamanishi K (ed), Human Herpesviruses: Biology, Therapy, and Immunoprophylaxis, Cambridge.

4. Sinclair JH, Reeves MB. 2013. Human cytomegalovirus manipulation of latently infected cells. Viruses 5:2803–2824.

5. Frenkel LD. 2016. Comparing Congenital Zika and Cytomegalovirus Affliction. Pediatr Infect Dis J 35:1371–1372.

6. Yang R, Zhang R, Zhang Y, Huang Y, Liang H, Gui G, Gong S, Wang H, Xu M, Fan J. 2019. Risk Factors Analysis for Human Cytomegalovirus Viremia in Donor+/Recipient+ Hematopoietic Stem Cell Transplantation. Lab Med doi:10.1093/labmed/lmz030.

7. Murphy E, Yu D, Grimwood J, Schmutz J, Dickson M, Jarvis MA, Hahn G, Nelson JA, Myers RM, Shenk TE. 2003. Coding potential of laboratory and clinical strains of human cytomegalovirus. Proc Natl Acad Sci U S A 100:14976–14981.

8. Murphy E, Rigoutsos I, Shibuya T, Shenk TE. 2003. Reevaluation of human cytomegalovirus coding potential. Proc Natl Acad Sci U S A 100:13585–13590.

9. Murthy S, O’Brien K, Agbor A, Angedakin S, Arandjelovic M, Ayimisin EA, Bailey E, Bergl RA, Brazzola G, Dieguez P, Eno-Nku M, Eshuis H, Fruth B, Gillespie TR, Ginath Y, Gray M, Herbinger I, Jones S, Kehoe L, Kuhl H, Kujirakwinja D, Lee K, Madinda NF, Mitamba G, Muhindo E, Nishuli R, Ormsby LJ, Petrzelkova KJ, Plumptre AJ, Robbins MM, Sommer V, Ter Heegde M, Todd A, Tokunda R, Wessling E, Jarvis MA, Leendertz FH, Ehlers B, Calvignac-Spencer S. 2019. Cytomegalovirus distribution and evolution in hominines. Virus Evol 5:vez015.

10. Engel P, Angulo A. 2012. Viral immunomodulatory proteins: usurping host genes as a survival strategy. Adv Exp Med Biol 738:256–276.

11. Hensel GM, Meyer HH, Buchmann I, Pommerehne D, Schmolke S, Plachter B, Radsak K, Kern HF. 1996. Intracellular localization and expression of the human cytomegalovirus matrix phosphoprotein pp71 (ppUL82): evidence for its translocation into the nucleus. J Gen Virol 77 (Pt 12):3087–3097.

12. Dunn W, Chou C, Li H, Hai R, Patterson D, Stolc V, Zhu H, Liu F. 2003. Functional profiling of a human cytomegalovirus genome. Proc Natl Acad Sci U S A 100:14223–14228.

13. Yu D, Silva MC, Shenk T. 2003. Functional map of human cytomegalovirus AD169 defined by global mutational analysis. Proc Natl Acad Sci U S A 100:12396–12401.

14. Kalejta RF, Shenk T. 2003. Proteasome-dependent, ubiquitin-independent degradation of the Rb family of tumor suppressors by the human cytomegalovirus pp71 protein. Proc Natl Acad Sci U S A 100:3263–3268.

15. Kalejta RF, Shenk T. 2003. The human cytomegalovirus UL82 gene product (pp71) accelerates progression through the G1 phase of the cell cycle. J Virol 77:3451–3459.

16. Saffert RT, Kalejta RF. 2007. Human cytomegalovirus gene expression is silenced by Daxx-mediated intrinsic immune defense in model latent infections established in vitro. J Virol 81:9109–9120.

17. Hwang J, Kalejta RF. 2007. Proteasome-dependent, ubiquitin-independent degradation of Daxx by the viral pp71 protein in human cytomegalovirus-infected cells. Virology 367:334–338.

18. Stiegler P, De Luca A, Bagella L, Giordano A. 1998. The COOH-terminal region of pRb2/p130 binds to histone deacetylase 1 (HDAC1), enhancing transcriptional repression of the E2F-dependent cyclin A promoter. Cancer Res 58:5049–5052.

19. Yang X, Khosravi-Far R, Chang HY, Baltimore D. 1997. Daxx, a novel Fas-binding protein that activates JNK and apoptosis. Cell 89:1067–1076.

20. Trgovcich J, Cebulla C, Zimmerman P, Sedmak DD. 2006. Human cytomegalovirus protein pp71 disrupts major histocompatibility complex class I cell surface expression. J Virol 80:951–963.

21. Fu YZ, Su S, Gao YQ, Wang PP, Huang ZF, Hu MM, Luo WW, Li S, Luo MH, Wang YY, Shu HB. 2017. Human Cytomegalovirus Tegument Protein UL82 Inhibits STING-Mediated Signaling to Evade Antiviral Immunity. Cell Host Microbe 21:231–243.

22. Burdette DL, Monroe KM, Sotelo-Troha K, Iwig JS, Eckert B, Hyodo M, Hayakawa Y, Vance RE. 2011. STING is a direct innate immune sensor of cyclic di-GMP. Nature 478:515–518.

23. Ishikawa H, Barber GN. 2008. STING is an endoplasmic reticulum adaptor that facilitates innate immune signalling. Nature 455:674–678.

24. Stamler JS, Simon DI, Osborne JA, Mullins ME, Jaraki O, Michel T, Singel DJ, Loscalzo J. 1992. S-nitrosylation of proteins with nitric oxide: synthesis and characterization of biologically active compounds. Proc Natl Acad Sci U S A 89:444–448.

25. Gould N, Doulias PT, Tenopoulou M, Raju K, Ischiropoulos H. 2013. Regulation of protein function and signaling by reversible cysteine S-nitrosylation. J Biol Chem 288:26473–26479.

26. Lindermayr C, Sell S, Durner J. 2008. Generation and detection of S-nitrosothiols. Methods Mol Biol 476:217–229.

27. Hao G, Derakhshan B, Shi L, Campagne F, Gross SS. 2006. SNOSID, a proteomic method for identification of cysteine S-nitrosylation sites in complex protein mixtures. Proc Natl Acad Sci U S A 103:1012–1017.

28. Dighiero P, Reux I, Hauw JJ, Fillet AM, Courtois Y, Goureau O. 1994. Expression of inducible nitric oxide synthase in cytomegalovirus-infected glial cells of retinas from AIDS patients. Neurosci Lett 166:31–34.

29. Nukui M, O’Connor CM, Murphy EA. 2018. The Natural Flavonoid Compound Deguelin Inhibits HCMV Lytic Replication within Fibroblasts. Viruses 10.

30. Eng JK, McCormack AL, Yates JR. 1994. An approach to correlate tandem mass spectral data of peptides with amino acid sequences in a protein database. J Am Soc Mass Spectrom 5:976–989.

31. Warming S, Costantino N, Court DL, Jenkins NA, Copeland NG. 2005. Simple and highly efficient BAC recombineering using galK selection. Nucleic Acids Res 33:e36.

32. Krishna BA, Miller WE, O’Connor CM. 2018. US28: HCMV’s Swiss Army Knife. Viruses 10.

33. Baldick CJ, Jr., Marchini A, Patterson CE, Shenk T. 1997. Human cytomegalovirus tegument protein pp71 (ppUL82) enhances the infectivity of viral DNA and accelerates the infectious cycle. J Virol 71:4400–4408.

34. Ma Z, Jacobs SR, West JA, Stopford C, Zhang Z, Davis Z, Barber GN, Glaunsinger BA, Dittmer DP, Damania B. 2015. Modulation of the cGAS-STING DNA sensing pathway by gammaherpesviruses. Proc Natl Acad Sci U S A 112:E4306–4315.

35. Wang X, Majumdar T, Kessler P, Ozhegov E, Zhang Y, Chattopadhyay S, Barik S, Sen GC. 2017. STING Requires the Adaptor TRIF to Trigger Innate Immune Responses to Microbial Infection. Cell Host Microbe 21:788.

36. Nevels M, Brune W, Shenk T. 2004. SUMOylation of the human cytomegalovirus 72-kilodalton IE1 protein facilitates expression of the 86-kilodalton IE2 protein and promotes viral replication. J Virol 78:7803–7812.

37. Bechtel JT, Shenk T. 2002. Human cytomegalovirus UL47 tegument protein functions after entry and before immediate-early gene expression. J Virol 76:1043–1050.

38. Sinzger C, Hahn G, Digel M, Katona R, Sampaio KL, Messerle M, Hengel H, Koszinowski U, Brune W, Adler B. 2008. Cloning and sequencing of a highly productive, endotheliotropic virus strain derived from human cytomegalovirus TB40/E. J Gen Virol 89:359–368.

39. Jaffrey SR, Snyder SH. 2001. The biotin switch method for the detection of S-nitrosylated proteins. Sci STKE 2001:pl1.

40. Bresnahan WA, Shenk TE. 2000. UL82 virion protein activates expression of immediate early viral genes in human cytomegalovirus-infected cells. Proc Natl Acad Sci U S A 97:14506–14511.

41. Lau L, Gray EE, Brunette RL, Stetson DB. 2015. DNA tumor virus oncogenes antagonize the cGAS-STING DNA-sensing pathway. Science 350:568–571.

42. Sun L, Wu J, Du F, Chen X, Chen ZJ. 2013. Cyclic GMP-AMP synthase is a cytosolic DNA sensor that activates the type I interferon pathway. Science 339:786–791.

43. Das S, Pellett PE. 2011. Spatial relationships between markers for secretory and endosomal machinery in human cytomegalovirus-infected cells versus those in uninfected cells. J Virol 85:5864–5879.

44. Diner EJ, Burdette DL, Wilson SC, Monroe KM, Kellenberger CA, Hyodo M, Hayakawa Y, Hammond MC, Vance RE. 2013. The innate immune DNA sensor cGAS produces a noncanonical cyclic dinucleotide that activates human STING. Cell Rep 3:1355–1361.

45. Chambers J, Angulo A, Amaratunga D, Guo H, Jiang Y, Wan JS, Bittner A, Frueh K, Jackson MR, Peterson PA, Erlander MG, Ghazal P. 1999. DNA microarrays of the complex human cytomegalovirus genome: profiling kinetic class with drug sensitivity of viral gene expression. J Virol 73:5757–5766.

46. Browne EP, Shenk T. 2003. Human cytomegalovirus UL83-coded pp65 virion protein inhibits antiviral gene expression in infected cells. Proc Natl Acad Sci U S A 100:11439–11444.

47. Mathers C, Schafer X, Martinez-Sobrido L, Munger J. 2014. The human cytomegalovirus UL26 protein antagonizes NF-kappaB activation. J Virol 88:14289–14300.

48. Barrett DM, Black SM, Todor H, Schmidt-Ullrich RK, Dawson KS, Mikkelsen RB. 2005. Inhibition of protein-tyrosine phosphatases by mild oxidative stresses is dependent on S-nitrosylation. J Biol Chem 280:14453–14461.

49. Kalejta RF, Bechtel JT, Shenk T. 2003. Human cytomegalovirus pp71 stimulates cell cycle progression by inducing the proteasome-dependent degradation of the retinoblastoma family of tumor suppressors. Mol Cell Biol 23:1885–1895.

50. Jia J, Arif A, Terenzi F, Willard B, Plow EF, Hazen SL, Fox PL. 2014. Target-selective protein S-nitrosylation by sequence motif recognition. Cell 159:623–634.

51. Jia J, Arif A, Willard B, Smith JD, Stuehr DJ, Hazen SL, Fox PL. 2012. Protection of extraribosomal RPL13a by GAPDH and dysregulation by S-nitrosylation. Mol Cell 47:656–663.

52. Guerra D, Ballard K, Truebridge I, Vierling E. 2016. S-Nitrosation of Conserved Cysteines Modulates Activity and Stability of S-Nitrosoglutathione Reductase (GSNOR). Biochemistry 55:2452–2464.

53. Kalejta RF. 2004. Human cytomegalovirus pp71: a new viral tool to probe the mechanisms of cell cycle progression and oncogenesis controlled by the retinoblastoma family of tumor suppressors. J Cell Biochem 93:37–45.

